# There and back again: when and how the world’s richest snake family (Dipsadidae) dispersed and speciated across the Neotropical region

**DOI:** 10.1101/2023.04.15.535132

**Authors:** Filipe C. Serrano, Matheus Pontes-Nogueira, Ricardo J. Sawaya, Laura R.V. Alencar, Cristiano C. Nogueira, Felipe G. Grazziotin

## Abstract

**Aim:** The widespread megadiverse Neotropical snake family Dipsadidae occurs in a large range of diverse habitats. Thus it represents an excellent model to study the diversification of Neotropical biota. Herein, by generating a time-calibrated species-level phylogeny, we investigate the origin and historical biogeography of Dipsadidae and test if its two main Neotropical subfamilies, Xenodontinae and Dipsadinae, have different geographical origins.

**Location:** Neotropical region.

**Taxon:** Dipsadidae (Serpentes).

**Methods:** We generated a new Bayesian time-calibrated phylogeny including sequences from six genes for 344 species, including 287 species of Dipsadidae. We subsequently estimated ancestral areas of distribution by comparing models in BioGeoBEARS: DEC (subset sympatry, narrow vicariance), DIVALIKE (narrow and wide vicariance), BAYAREALIKE (no vicariance and widespread sympatry), also testing jump dispersal.

**Results:** The best models show that Dipsadidae likely originated approximately 50 million years ago (mya) in Asia. Dispersal was a fundamental process in its historical biogeography. The DEC model with jump dispersal indicated that this family underwent a range extension from Asia and posterior vicariance of North and Central America ancestors. Both Xenodontinae and Dipsadinae originated in Central America and dispersed to South America during Middle Eocene, but did so to different regions (cis and trans-Andean South America, respectively). Xenodontinae entered cis-Andean South America around 39 mya and jump dispersed to the West Indies around 33 mya, while Dipsadinae entered trans-Andean South America multiple times 20 – 38 mya.

**Main conclusions:** Our results show that Dipsadidae has an Asian origin and that the two main Neotropical subfamilies originated in Central America, later dispersing to South America in different time periods. The current biogeographical patterns of the family Dipsadidae, the most species-rich snake family in the world, have likely been shaped by complex evolutionary and geological processes such as Eocene land bridges, Andean uplift and the formation of the Panama isthmus.

## Introduction

The Neotropical realm is a climatically and geologically diverse biogeographical region, encompassing a wide range of habitats, from the lush rainforests of the Amazon and Central America to the snow-covered peaks of the Andes. This diversity of habitats is the result of a rich and complex paleogeographical history between and within two continental landmasses — Central and South America — and the associated island systems (e.g., Galapagos, West Indies; Clapperton, 1993; Pennington *et al*., 2004; Rull, 2011; Hughes, Pennington & Antonelli, 2013). Even though major geological events such as the Gondwana breakup and the formation of volcanic hotspots happened during the Mesozoic era (Jokat, Boebel, König & Meyer, 2003; Wilf, Cúneo, Escapa, Pol & Woodburne, 2013), many geomorphological events relevant to modern-day Neotropical region occurred in the Cenozoic. These include mountain uplift in Central America and the Andes, the formation of the West Indies island system, a potential short-lived land-bridge connecting South America to the West Indies (the Greater Antilles and Aves Ridge, GAARlandia; Iturralde-Vinent & MacPhee, 1999; but see Ali & Hedges, 2021) and formation of the Isthmus of Panama, a contiguous landmass connecting Central and South America whilst separating the Atlantic and Pacific oceans (Graham 2009; Hoorn *et al*. 2010).

These geomorphological events and their abiotic and biotic consequences widely shaped the evolutionary history of the Neotropical biota, contributing for the Neotropics to be today the world’s most biodiverse region (Antonelli & Sanmartin 2011; Rull, 2011). Therefore, Neotropical faunal assemblages reflect several distinct biogeographical histories. While some clades likely originated by mid-Cretaceous vicariant event between South America and Africa (e.g., boid snakes: Noonan & Chippindale, 2006; Iguanian and Scleroglossan lizards: Albino & Brizuela, 2014), others later overwater dispersed from Africa (e.g., Epictine threadsnakes: Adalsteinsson, Branch, Trape, Vitt & Hedges, 2009; Platyrrhine monkeys and Caviomorph rodents: Defler 2019; South American Amphisbaenidae: Graboski, Grazziotin, Mott & Rodrigues, 2022) or from Asia, via North America (viperid snakes: Wüster, Peppin, Pook & Walker, 2008; turtles: Lichtig, Jasinski & Lucas, 2019). Furthermore, the more recent Great American Biotic Interchange (GABI) promoted dispersal and faunal admixture between Central and South American fauna — mainly mammals and birds (Bacon *et al*., 2015; Defler 2019; South American Amphisbaenidae: Graboski *et al*., 2022) — despite some evidence of pre-GABI dispersal (Heinicke, Duellman & Hedges, 2007; Agnolin, Chimento & Lucero, 2019). Other groups, such as reptiles, are thought to have been less directly involved in GABI, mostly diversifying in Central America with later dispersal to South America with few groups doing the reverse path (Vanzolini & Heyer 1985).

Widely distributed taxa represent ideal models to study biogeographic processes in the Neotropics (Colston *et al*., 2013; Torres-Carvajal, Echevarría, Lobos, Venegas & Kok, 2019; Azevedo *et al*., 2020). Snakes are exceptionally diverse in the Neotropical realm, where roughly one-third of all species occur (Guedes *et al*., 2017; Roll *et al*., 2017; Nogueira *et al*., 2019). Dipsadidae (Bonaparte, 1838) is the richest snake family in the Neotropics with over 700 known species, which are diverse in diet, habitat use, and morphology (Cadle & Greene, 1993; Serrano *et al*., in prep.). This high level of biological variation among dipsadids is reflected on the distributional patterns and the phylogenetic relationships within the family, making it a promising but scarcely explored model to evaluate biogeographic hypotheses of diversification (Grazziotin *et al*. 2012, Zaher *et al*., 2019). It comprises four well-known groups: the monophyletic and highly diverse subfamilies Dipsadinae and Xenodontinae, which are widespread in the Neotropical realm (Cadle & Greene, 1993); plus two relict groups, one distributed in North America that includes the subfamily Carphophiinae and the genera *Heterodon* and *Farancia* (Pinou, Vicario, Marschner & Caccone, 2004), and another exclusively distributed in Asia composed of the genera *Thermophis* and *Stichophanes* (Huang, Liu, Guo, Zhang, & Zhao; Grazziotin *et al*., 2012; Zaher *et al*., 2019).

Despite the uncertainty around the family’s geographical origin, hypotheses of ancestral distribution have ranged from a Gondwanan distribution (Cadle, 1985), an Asian origin followed by a dispersal from Asia via North America (Cadle, 1985) and an African origin followed by a trans-Atlantic dispersal to South America (Cadle, 1984), possibly followed by a dispersal to North America (Duellman, 1979). Recent phylogenetic studies have supported an Asian-North American dispersal event based on the interpretation of the successive sister-group relationship between the Asian genera *Thermophis* and *Stichophanes* and the clade composed by American dipsadids (Grazziotin *et al*. 2012; Zaher *et al*., 2019). This Asian-North American dispersal event has been supposed even before the studies positioning of *Thermophis* and *Stichophanes* (Cadle, 1985), and it is frequently associated with the formation of the Beringian Bridge during the Miocene, around 16–10 mya. The same hypothesis is presented as the general biogeographical explanation for the presence of other snake families, such as Colubridae and Natricidae in the New World (Vidal, Dewynter & Gower, 2000; Pinou *et al*., 2004).

However, in recent studies, the estimated divergence between American and Asian dipsadids is older than the Miocene. Zaher *et al*. (2018; 2019) estimated this divergence between 22 mya and 27 mya, around the transition between the Oligocene and Miocene. Other studies have suggested older dates, pointing to a divergence between Asian and American dipsadids dated in the transition between the Eocene and Oligocene (between 26 and 36 mya; Entiauspe-Neto, *et al*. *in press*). An alternative hypothesis supporting pre-Miocene divergence times is related to cladogenic events as the opening of the Greenland corridor approximately 48 mya.

Within the diversity of dipsadids, some studies restricted to small groups of species (e.g., *Leptodeira*: Daza, Smith, Páez & Parkinson, 2009; Imantodini: Mulcahy, 2007; *Thermophis*: Huang *et al*., 2009) have only reconstructed recent biogeographical patterns and attained some estimates of divergence times but achieved inconclusive results regarding the ancestral range distribution and biogeographic processes of the main groups. Since the classical studies of Cadle (1984a, 1984b, 1984c), the evolutionary history of the two major dipsadid subfamilies has been understood as reflecting independent origins and processes of diversification. Following Cadle’s hypothesis, Dipsadinae originated in Central America, where the subfamily diversified and further dispersed to South America. Xenodontinae, on the other hand, would have originated and diversified in South America, and from there, dispersed to Central America. Although Duellman (1979) suggested a different scenario—a common South American origin for Dipsadidae and further dispersal to Central and North America — the hypothesis provided by Cadle was well accepted by the herpetological community, and it has been supported by further studies (Cadle & Greene, 1993; Vidal et al., 2000; Zaher et al., 2009; Hedges, Couloux & Vidal, 2009; Vidal et al., 2010; Grazziotin et al., 2012; Zaher et al., 2018; Zaher et al., 2019). Cadle also suggested that the divergence between both subfamilies had happened during the late Palaeocene–Eocene separation of Central and South America, around 40–60 mya (Cadle, 1985). However, recent studies have estimated divergence times between Dipsadinae and Xenodontinae varying around 19 mya and 24 mya, during the Late Miocene (Zaher *et al*. 2018; 2019).

Zaher *et al*. (2019) also suggested that the sister group affinities retrieved between Diaphorolepidini (an exclusive South American tribe) and the remaining Dipsadinae, on the one hand, and Conophiini (an exclusive Central American tribe) and the remaining Xenodontinae, on the other hand, points to a complex historical scenario of origin and diversification of the two main Central- and South-American dipsadid lineages than previously thought (Cadle, 1985; Cadle & Greene, 1993). Therefore, both the family’s origin and its overall biogeographical history, such as timing and route of dispersal between Central and South America, remains uncertain.

Historical biogeography (Posadas, Crisci & Katinas, 2006) is an essential tool to understand the origin and composition of current Neotropical biotas such as snake assemblages since biogeographical processes such as dispersal, vicariance, and extinction strongly influence local and regional biodiversity through time (Ricklefs, 1987; Moritz, Patton, Schneider & Smith, 2000; Crisci, 2001). However, comprehensive studies on Neotropical historical biogeography have been severely hampered by the lack of detailed phylogenetic hypotheses and distributional data (Bagley & Johnson, 2014) as well as analytical limitations (Landis, Matzke, Moore & Huelsenbeck, 2013; Matzke, 2013). Despite information available on the distribution, richness and phylogenetics of diverse groups such as snakes being increasingly available (López-Aguirre, Hand, Laffan & Archer, 2018; Nogueira *et al*., 2019; Azevedo *et al*., 2020), their historical biogeography is complex and still poorly understood.

Here, we generate and use a comprehensive time-calibrated phylogeny and a Bayesian estimation of the ancestral geographical ranges aiming to: (1) infer the most likely distribution of ancestral lineages of Dipsadidae, (2) reconstruct the historical biogeography of dipsadid snakes in the Neotropical region; and (3) complement the current knowledge of paleogeographical scenarios related to the diversification and current patterns of distribution of dipsadids in Central and South America. Specifically, we tested the hypotheses that: i) Dipsadidae had an Asian origin with dispersal via North America; and ii) Dipsadinae and Xenodontinae — the two Neotropical subfamilies — have different geographical origins (Central and South American, respectively).

## Materials and Methods

### Phylogenetic tree

We based our Bayesian phylogenetic analysis on the molecular dataset from Zaher *et al*. (2018), the most complete and up-to-date available dataset considering the diversity of Dipsadidae. The concatenated matrix included DNA sequences of six genes (12S, 16S, cytb, bdnf, c-mos, and nt-3) for 344 species representing the families Dipsadidae, Pseudoxenodontidae, Colubridae, Calamariidae, Sibynophiidae, Grayiidae, Natricidae, Viperidae, Pareidae, and the superfamily Elapoidea. The boids *Eryx conicus* and *Boa constrictor* were included to root the phylogenetic tree. The dataset is largely biased towards Dipsadidae (287 species, 83.4% of species in the phylogeny), with 283 New World species (84 genera), of which 10 (five genera) belong to the subfamily Carphophiinae, 167 (54 genera) to Xenodontinae, and 106 (23 genera) to Dipsadinae. The Asian incertae sedis Dipsadidae genera *Thermophis* (three species) and *Sticophanes* (one species) are also included in the molecular dataset to allow the estimation of the origin and early evolution of South American dipsadids. Overall, our sample of Dipsadidae represents nearly a third of all valid species for this family (Uetz *et al*., 2020).

To determine the optimal partitioning scheme and nucleotide substitution models of DNA, we used PartitionFinder v2.1.1 (Lanfear, Calcott, Ho & Guindon., 2012). We previously partitioned our concatenated matrix based on gene fragments and we tested all models implemented in MrBayes 3.1.2 (Ronquist & Huelsenbeck, 2003) through Bayesian Information Criterion (BIC), while using the ‘greedy’ algorithm (Lanfear, 2012).

Although Zaher *et al*. (2018) have phylogenetically analyzed the same matrix using Maximum Likelihood (ML), they only estimated divergence times based on a considerably reduced matrix (with 67 terminals), focusing on the evolution of Pseudalsophis in the Galápagos Archipelago. Here, we greatly extended their analysis by performing a time-calibrated Bayesian inference in MrBayes 3.1.2 using the complete matrix with 344 terminals, aiming to estimate divergence time within the whole diversity of Dipsadidae. We defined a set of topological constraints based on the ML topology presented by Zaher *et al*. (2018) to reduce the tree space and decrease the running time of our analysis. The set of topological constraints is listed in the nexus file (available at figshare [figshare address]). Node calibration points were defined based on the fossil record and we used similar ages and fossil interpretations as described by Zaher *et al*. (2018) and Zaher *et al*. (2019). The list of calibration points and their respective references are available in the Supp. Mat. S1).

We set the branch length prior as a birth-death clock model (Yang and Rannala, 1997), with speciation and extinction probabilities set to exponential (lambda = 10) and beta (alpha = 1 and beta = 1) distributions, respectively. We divided the total number of terminals in our molecular matrix by the approximate total number of extant alethinophidians (Uetz *et al*., 2020) and we set the sample probability to 0.109. For the model of variation of the clock rate across lineages, we used the independent gamma rates (IGR) model (Ronquist *et al*. 2012) with the parameter IGRvar — the amount of rate variance across branches — set to the exponential (lambda = 10). To set the clock rate, we followed Pyron (2017), and we used a log normal distribution with a mean corresponding to the log of the average number of substitutions per site from root to tips estimated from the tree provided by Zaher *et al*. (2018), divided by the mean root age (-3.295561). The standard deviation for the log normal distribution was set as the exponent of the mean (1.037742).

We implemented this analysis in two independent runs with eight Markov Chains Monte Carlo (MCMC, one cold and seven incrementally heated) and 50 million generations. To generate the 50% majority rule consensus tree, a conservative burn-in of 25% was applied after checking the log-likelihood scores and the split-frequencies of the runs, and all sampled trees prior to reaching these generations were discarded.

Clades with support values ≥ 0.85 were considered well-supported. We combined the resulting trees from the two runs using the sumt command in MrBayes, and eventual polytomies were randomly solved by adding small branch-lengths (0.0001) using functions from the ‘ape’ package (Paradis & Schliep, 2019) in R 3.5.2 (R Core Team, 2019). The complete time-calibrated Bayesian tree was pruned to Dipsadidae and its closest sister group to implement further historical biogeographical analysis.

### Biogeographical analysis and ancestral range estimation

Several methods have been used to define biogeographical units (Ferrari, 2017), variating from areas of endemism (Morrone, 1994; Linder, 2001), biotic elements (Hausdorf & Hennig, 2003), bioregions (Edler, Guedes, Zizka, Rosvall & Antonelli, 2017), tectonic plates (Sanmartín & Ronquist, 2004) and zoogeographic realms (Holt *et al*., 2013), among others. All of them present strengths and weakness (Ferrari 2017), but regarding the scope of our study, and the geographical and phylogenetic pattern of Dipsadidae, we based our units combining a modified version of zoogeographic realms (Holt *et al*. 2013) with areas of endemism (Morrone, 2010; Morrone *et al*., 2014). We used the ‘Mexican transition zone’ as the limit between North and Central America since it separates the Nearctic and Neotropical regions (Morrone, 2010; Morrone, Escalante & Rodriguez-Tapia, 2017), and northern Nicaragua as the limit to Central America because it represents the austral border of the Mesoamerican Dominion (Morrone *et al*., 2014) and its southern portion (Panama, Costa Rica and southern Nicaragua) is much younger than its northern portion due to their different geological histories (Bacon *et al*., 2015; O’Dea *et al*., 2016). Since we aimed to understand only major biotic exchanges between insular and continental landmasses, the West Indies were treated as a single area to decrease the number of biogeographical units and consequently the models’ running time. These biogeographic units are congruent with several main geographic/phylogenetic groups of Dipsadidae included in our analysis (Asian dipsadids: *Thermophis* and *Stichophanes*; North American dipsadids: Carphophiinae, Heterodon and Farancia; Central American dipsadids: most of the Dipsadinae; cis-Andean South American dipsadids: most of the Xenodontinae; trans-Andean South American dipsadids: rare radiations present in both subfamilies, Xenodontinae and Dipsadinae; and West Indian dipsadids: mainly the tribe Alsophiini).

We considered six biogeographical units (Fig. 1a), assigning each species distribution to one or more than one of them: (A) Asia, (B) North America (American continent north of the Trans-Mexican Volcanic Belt), (C) Central America (from the Trans-Mexican Volcanic Belt to northern Nicaragua), (D) the West Indies, (E) Trans-Andean South America (from western slopes of the Andes to the Pacific Ocean shores) and (F) cis-Andean South America (from eastern slopes of the Andes to the Atlantic Ocean shores). We constrained the maximum number of occupied units to three, since none of the extant species occurs in more than three areas.

**Figure 1.**
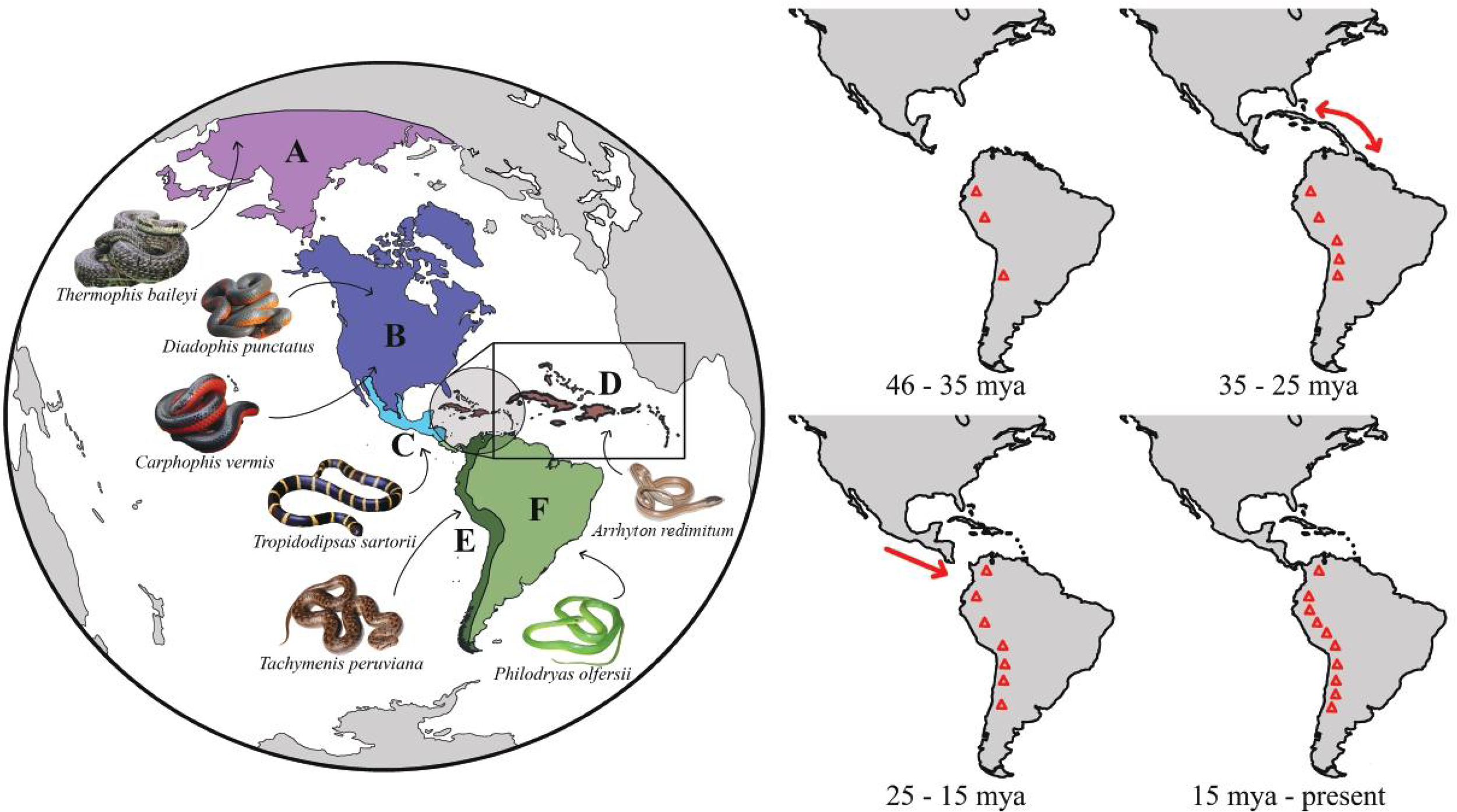
a) Biogeographical units considered in this study and their representative species; A – Asia, B – North America, C – Central America, D – West Indies, E – trans-Andean South America and F – cis-Andean South America. b) Relevant geomorphological events in the Neotropical region since the Eocene Epoch (56 to 33.9 million years ago – mya). Red arrows represent land connections and red triangles represent increasing elevation in the Andes.

We estimated the ancestral ranges for Dipsadidae using the *BioGeoBEARS* package (Matzke, 2013) in R 3.5.2 (R Core Team, 2019), using variations of the likelihood models DEC (Dispersal-Extinction-Cladogenesis; Ree & Smith, 2008), DIVA-like (Dispersal-Vicariance Analysis; Ronquist, 1997) and BayArea-like (Bayesian Inference of Historical Biogeography for Discrete Areas; Landis *et al*., 2013). The DEC (Dispersal-Extinction Cladogenesis — Ree & Smith, 2008; Matzke, 2013) model assumes that derived lineages following cladogenesis can only inherit a single range area, which is a subset of their ancestor’s range; DIVAlike (Ronquist & Sanmartin, 2011) which allows vicariant events, but does not allow for sympatric-subset speciation by derived lineages. BAYAREAlike (Landis *et al*., 2013), on the other hand, assumes that no range evolution occurs at cladogenesis, and derived lineages inherit the same range of the ancestral state, making it a heavily dispersalist model.

Although we tested all models implemented in *BioGeoBEARS*, we acknowledge that statistical comparison among models without incorporating subjective biological knowledge can favour models that, despite increasing the data likelihood, do not necessarily incorporate the most probable historical scenario (Sanmartín, 2021). We assume that for an old (probably more than 40 my old) wide dispersed taxa (four continents) like Dipsadidae, evolution by vicariance needs to be considered in biogeographical models, even if it occurs at a low rate. Therefore, we maintained BAYAREAlike models in our analysis only to test the relative importance of scenarios mainly driven by dispersal (see results below), but we base our main discussion on the best models that allow vicariant processes.

We furthermore compared the above models with the added +j parameter, which allows founder-event speciation and was added due to its potential importance in reconstructing insular historical biogeography (Klaus and Matzke; 2020; Matzke, 2022; but see Ree and Sanmartín; 2018). To each model we also added a time-stratified matrix with dispersal probabilities (Supp. Mat. 2) between pairs of areas specified based on geological events occurring in each period (Fig. 1b), varying between 0.1 (unlikely), 0.5 (probable) and 1 (likely). For this matrix we considered potentially relevant events (Figure 1b) at 46 mya [million years ago] (origin of the clade), 35 mya (potential uplift of GAARlandia or stepping stone islands; Iturralde-Vinent & MacPhee, 1999), 30 mya (disappearance of GAARlandia or stepping stone islands; Iturralde-Vinent & MacPhee, 1999), 25 mya (approximation of the Central American and South American tectonic plates; Montes *et al*., 2012) and 15 mya (complete formation of the Panama Ishtmus; Bacon *et al*., 2015 but see O’Dea *et al*., 2016). All models were implemented in the Maximum Likelihood framework of BioGeoBEARS, (Matzke, 2013). In total, we implemented six Maximum Likelihood models which were compared via Akaike information Criterion – AIC (Akaike, 1974; Wang, 2006).

## Results

### Phylogeny and divergence time estimation

Our phylogeny (Fig. 2) suggests a crown age of Colubroidea of 56.6 my (49.2-63.7 my 95% HPD), with the main split between Dipsadidae — strongly supported as monophyletic — and the remaining Colubroidea occurring in mid Eocene approximately 49.1 mya (44.1-55.4 mya 95% HPD) (available at 10.6084/m9.figshare.22634854). The split between Asian and American Dipsadidae occurred at 44.9 mya (40.1-50.2 mya 95% HPD), with the more species-rich Neotropical Dipsadidae splitting from the North American relictual clade at 43.1 mya (38.2 -47.3 mya 95% HPD). Both Xenodontinae and Dipsadinae were strongly recovered as monophyletic, while Carphophiinae was recovered as non-monophyletic. While most clades within Xenodontinae were well resolved (bar the *Erythrolamprus* and *Helicops* genera and the Tachymenini tribe, for instance), several clades within Dipsadinae showed low to moderate support with the most noticeable being the Dipsadini tribe and the *Atractus* + *Geophis* clade.

**Figure 2.**
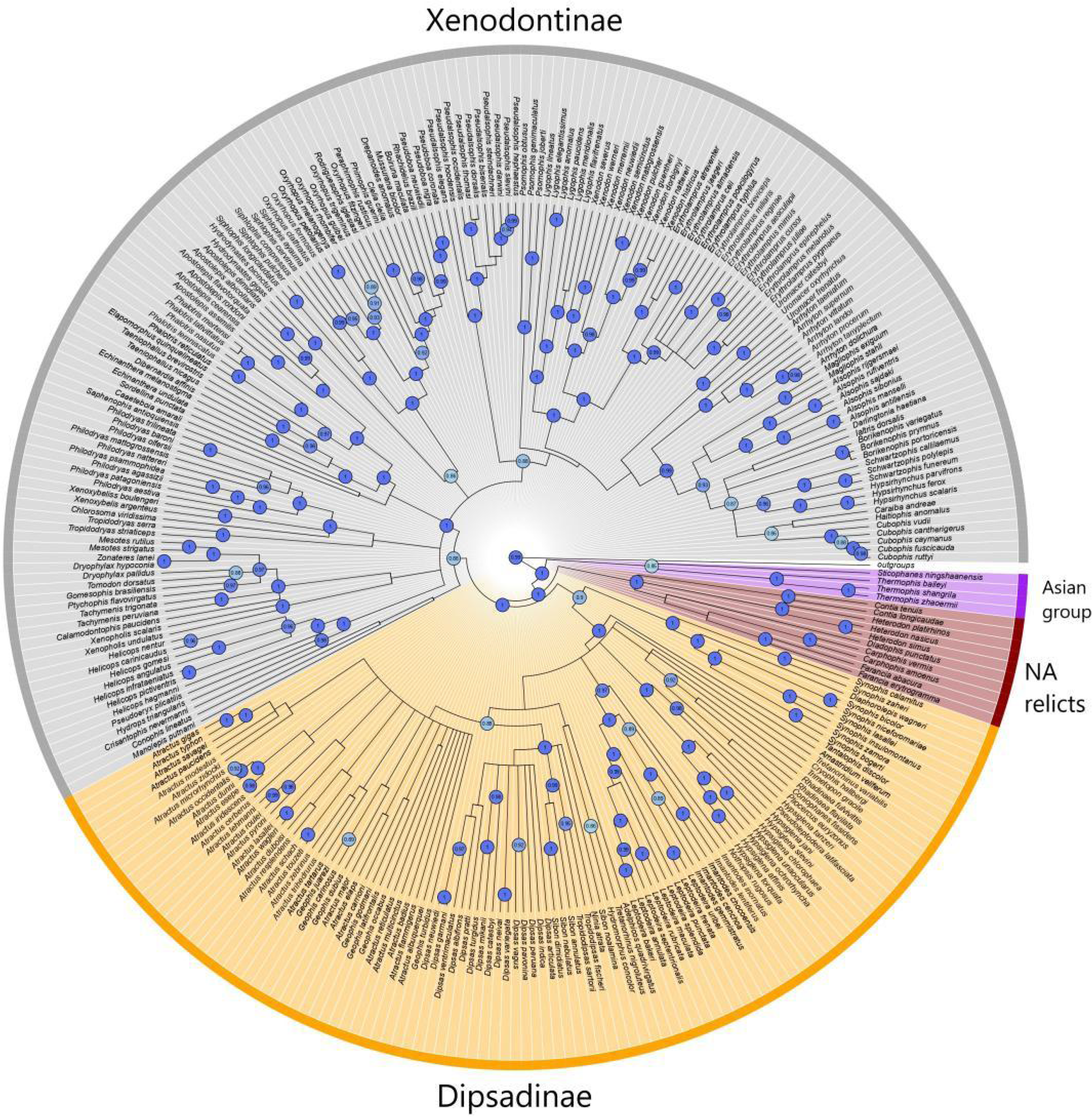
Time-calibrated Bayesian consensus phylogeny of Dipsadidae, with major groups represented: Xenodontinae (grey), Dipsadinae (orange), Carphophiinae (North American relicts, dark red) and Asian dipsadids (purple). Blue circles indicate statistical support for nodes > 85%

### Ancestral range estimation

The best fitted model was BAYAREALIKE +j (AICc = 619.9), followed by DEC +j (AICc = 647.2). This highlights the importance of dispersal for this snake clade, especially since the founder-event parameter was present in the three best models (Table 1). It also highlights that the anagenetic processes and range heritage were more important in the evolution of the dipsadids than the cladogenetic processes. However, as stated before, since BAYAREALIKE does not consider vicariant processes, we illustrate the historical biogeography of Dipsadidae with DEC +j. The most recent ancestor of Dipsadidae likely occurred in Asia, splitting from its sister group (the family Pseudoxenodontidae) during the Early Eocene. The clade’s extension of distribution to the New World (current North America and Central America) was then followed by a vicariant event between the Asian dipsadids and the American clade around 44.6 mya (40.1 – 50.2 mya 95% HPD) (Fig. 3). In the Mid Eocene, around 42.8 mya (37.6 – 48.6 mya 95% HPD), there was another vicariant event splitting the Carphophiinae subfamily in North America and the ancestor of the speciose Neotropical dipsadids in Central America. From then, around 42 mya, the two current major Neotropical subfamilies underwent distinct biogeographical processes. For Xenodontinae, a small lineage remained in Central America (Conophini), while the ancestor of the subfamilly dispersed into cis-Andean South America via jump dispersal. The ancestor lineage of Dipsadinae remained in Central America, with a further jump dispersal by the ancestor of the small lineage Diaphorolepidini to trans-Andean South America around the Eocene – Oligocene transition. Thereafter, Xenodontinae mainly maintained a cis-Andean distribution, except for the Alsophini clade, which underwent a major jump dispersal event to the West Indies during the early Oligocene, around 33.0 mya. The subfamily Dipsadinae, on the other hand, underwent many relevant biogeographical changes, especially since 30.7 mya, where the *Hypsiglena* + *Pseudoleptodeira* clade majorly reverted its distribution to North America. Compared to Xenodontinae, the occupation of South America by previously Central American dipsadines occurred much later, during the Oligo-Miocene transition, and by several jump dispersal events: at around 25.4 mya for the tribe Dipsadini and at around 22.3 mya for the genus Atractus. Overall, range extensions (e.g., range extension of a trans-Andean species to Central America) occurred at more recent times during Late Miocene and mainly within the subfamily Dipsadinae. Major events are summarized in Fig. 4.

**Table 1.** The best-fitted models of ancestral range estimation of Dipsadidae with BioGeoBEARS, all including a transition matrix. Model comparison based on log-likelihood (LnL), the corrected Akaike information criterion; n, number of parameters; d, rate of dispersal; e, rate of extinction; j, relative probability of founder-event speciation. The best model is shown in bold.

| Model | LnL | n | d | e | j | AICc |
| --- | --- | --- | --- | --- | --- | --- |
| **BAYAREALIKE +j** | **-306.9** | **3** | **0.0035** | **0.0003** | **0.039** | **619.9** |
| DEC +j | -320.6 | 3 | 0.0055 | 1.00E-12 | 0.03 | 647.2 |
| DIVALIKE +j | -332.6 | 3 | 0.006 | 1.00E-12 | 0.033 | 671.3 |
| DEC | -340.6 | 2 | 0.0079 | 1.00E-12 | 0 | 685.3 |
| DIVALIKE | -350.2 | 2 | 0.0095 | 1.00E-12 | 0 | 704.4 |
| BAYAREALIKE | -370.6 | 2 | 0.0046 | 0.016 | 0 | 745.2 |

**Figure 3.**
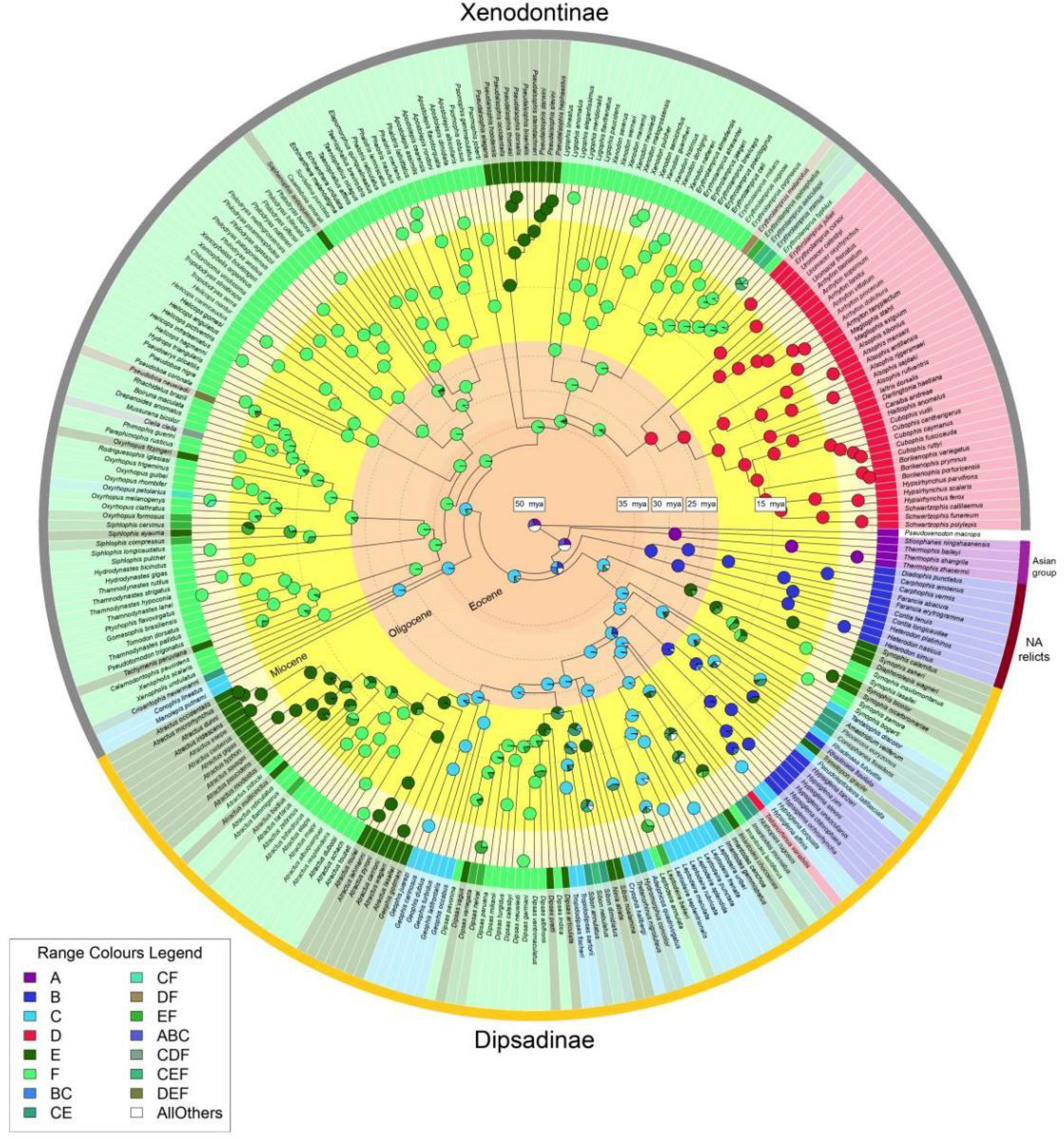
Ancestral area estimations from the DEC+j model implemented in BIOGEOBEARS. The most probable ancestral areas are mapped by pie charts at each node and the actual occurrence of each specie is colour coded next to the species name (see legend). Orange and yellow-ish circles inside the phylogeny indicate geological epochs (Miocene, Oligocene and Eocene are named). Dashed circles represent the time divisions present on the time-stratified matrix, with time in millions of years ago (mya) indicated at the white boxes.

**Fig. 4.**
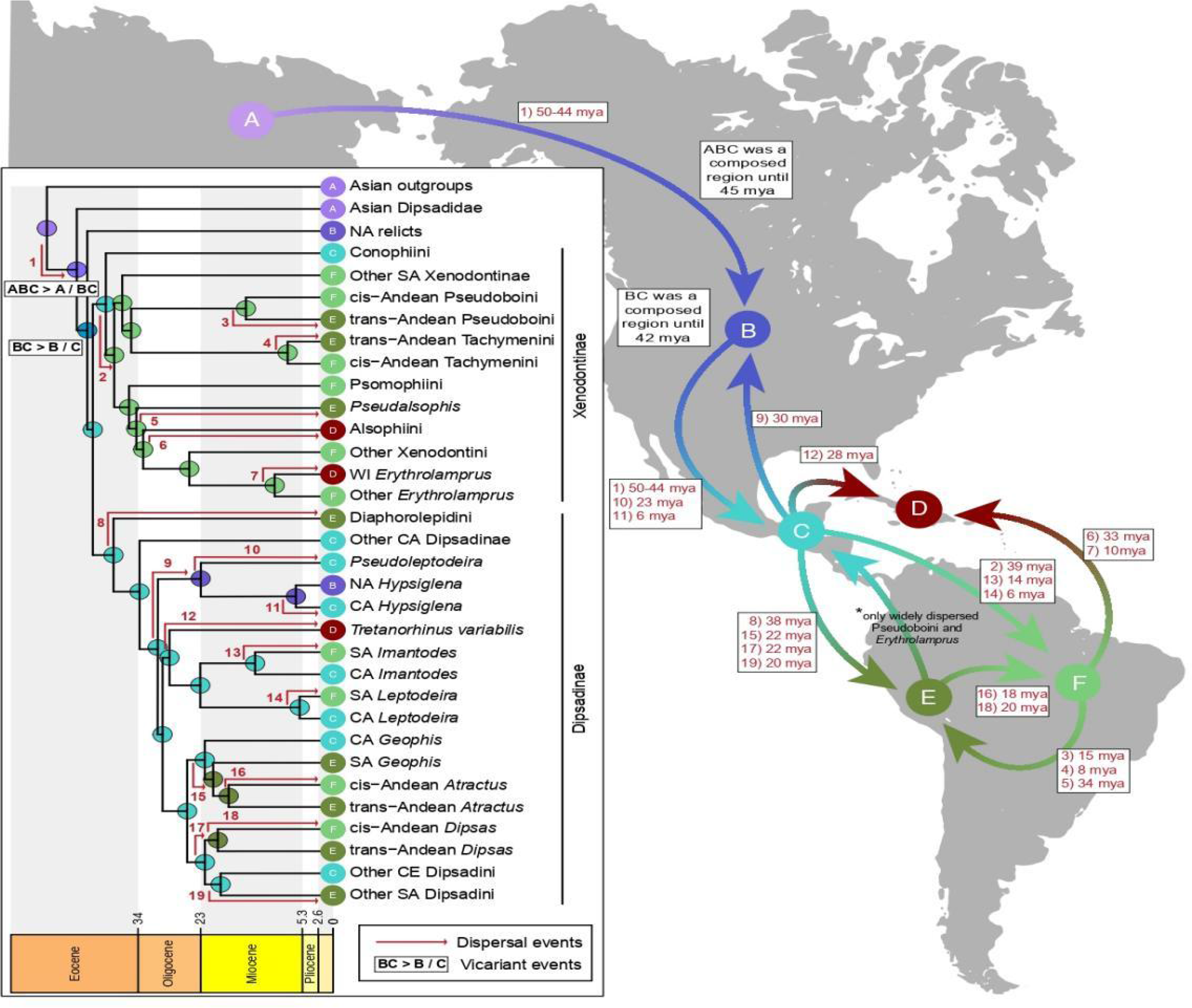
Summary of major biogeographical events of Dipsadidae. The purple circle represents the likely origin of the family, while arrows represent dispersal within the family at different time periods between regions. Inset: summarized phylogeny for representative taxa with numbered relevant dispersal and vicariant events.

## Discussion

Overall, we reconstruct the complex biogeographical history of the family Dipsadidae, the most species rich clade of Neotropical snakes and an important component of Neotropical biodiversity. Our results show that Dipsadidae has an Asian origin, corroborating our first hypothesis, and that the two main Neotropical subfamilies likely originated in Central America, contrary to our second hypothesis.

### Origin of New world dipsadids

Our findings strongly corroborate an Asian origin for dipsadids, as previously suggested (Cadle, 1984c; Grazziotin *et al*., 2012; Zaher *et al*., 2019), and thus challenge studies that suggested that the Dipsadidae could have an African or Gondwanan origin (Cadle, 1985) or that Dipsadidae could have dispersed from South America to North America during its early diversification (Duellman, 1979). This origin is also consistent with the current distribution of its closest clades, including the Asian Pseudoxenodontidae and Natricidae. In spite of being almost globally widespread (i.e. occurring in the Paleartic, Neartic and Afrotropical regions), Natricidae is mostly absent from the Neotropical region and its ancestral distribution is Asian (Deepak *et al*., 2022). This validates our first hypothesis of an Asian origin for the Dipsadidae and subsequent dispersal to North America, possibly via the Beringia Land Bridge. This land bridge is estimated to have connected the Palearctic and Nearctic realms during the Eocene (33-55 mya; Wolfe, 1975; Baskin & Baskin, 2016), being covered by warmer boreotropical forests which would have been suitable for ectotherms (Sanmartín, Enghoff & Ronquist, 2001; Townsend, Leavitt & Reeder, 2011; Baskin & Baskin, 2016; Graham, 2018). This dispersal pattern is coeval with other squamate taxa (Dibamid lizards: Townsend *et al*., 2011) and similar, albeit earlier than, coral snakes (Kelly, Barker, Villet & Broadley, 2009), lampropeltine rat snakes (Burbrink, Chen, Myers, Brandley & Pyron, 2012) and crotaline vipers (Wüster *et al*., 2008; Alencar *et al*., 2016). Alternatively, Dipsadidae could have reached North America from Asia via North Atlantic Land Bridges, especially the Thulean bridge, which were also present at the time of their origin (Tiffney, 1985; Jian *et al*., 2019). The Thulean land bridge connected southern Europe to Greenland, which in turn was connected to eastern North America and was available throughout the Early Tertiary until its submersion approximately 50 mya (Tiffney, 1985; Jian *et al*., 2019). Both plants and vertebrates have been suggested to have migrated via climatically suitable forest-covered North Atlantic Land Bridges (Sanmartín *et al*., 2001; Jian *et al*., 2019). However, dispersal via the Thulean bridge would imply that Dipsadidae once occupied and then went extinct in most of the Eurasian continent. While fossils associated with Dipsadidae (Paleoheterodon and Heterodon) have been described from southern Europe and North America, these are dated to Miocene/early Pliocene and could likely be a posterior incursion of North American fauna into Europe via the North Atlantic Greenland-Faroes bridge. Therefore, while both dispersal routes are possible, it likely that the geographically closer Beringia bridge likely provided a more suitable intercontinental dispersal route, as also suggested for other reptiles (e.g., Chen *et al*., 2013; Townsend *et al*., 2011).

### The distinct processes shaping the diversity of dipsadines and xenodontines

We show that the main cladogenetic event originating both Neotropical subfamilies of Dipsadidae (Dipsadinae and Xenodontinae) must have occurred in Central America, prior to their dispersal to South America, as hypothesized for different clades of the Neotropical herpetofauna (Vanzolini & Heyer, 1985). Thus, our results rejected the hypothesis of different geographical origins for Dipsadinae and Xenodontinae as suggested by Cadle & Greene (1993).

The DEC + j model shows that Dipsadidae has dispersed to South America several times during its diversification. Both subfamilies originated and begun to diversify in the Middle Eocene, when a major increase in temperature – the Middle Eocene Climatic Optimum or MECO – took place, which has been shown to have increase the diversity of plants and mammals (Woodburne *et al*., 2014; Fernandez, Santamarina, Palazzesi, Tellería, & Barreda, 2021). Numerous other significant intercontinental faunal dispersals have been documented for this period for many vertebrates (Beard, Qi, Dawson, Wang & Li, 1994; Chaimanee *et al*., 2012). Furthermore, both Neotropical subfamilies, despite first entering South America quasi-simultaneously around 40 mya, have different biogeographical histories, despite the common biogeographical origin. Xenodontinae likely incurred in a single colonization through jump dispersal to South America by a Central American ancestor in the Middle Eocene (∼ 39 mya), that was followed by quasi-isolation of the group in the region (Simpson, 1980; Cadle 1985). The exceptions to this isolation are dispersing species that returned to Central America and/or some lineages that dispersed to the West Indies, including the jump dispersal by the Alsophinii clade (Fig 3). The presence of Dipsadinae in South America is also explained by jump dispersal to South America from a Central American ancestors, although through multiple events (at least five) occured in the Middle Eocene (Diaphorolepidini), Early Miocene (South American species of goo-eaters), and in the Middle (South American species of *Imantodes*) and Late Miocene (South American species of *Leptodeira*). The time of the first dispersal event of dipsadines (∼ 38 mya) coincides with that estimated for Xenodontinae (Fig 3), although the time frame of these dispersal events from Central to South America indicated by our results is not congruent with paleogeographical reconstructions of a contiguous connection of the two continents, which suggested a large seaway separating the two landmasses (Montes *et al*., 2012, but see Coates & Stallard, 2016). Although this seaway likely represented a major obstacle to biotic interchange, the migration rate between the two continental masses has already been shown to have significantly increased around 41 mya (Bacon *et al*., 2015). Long-distance rafting and over-water dispersal from continental landmasses could explain such dispersal events (O’Dea *et al*., 2016), especially stepping-stone dispersal via islands in the present-day Caribbean Sea, as suggested for other species (ants: Archibald, Cover & Moreau, 2006; butterflies: Condamine, Silva-Brandão, Kergoat & Sperling, 2012; carnivorous plants: Ellison *et al*., 2012). Even though most islands of the West Indies were not above sea level before about 40 mya for Greater Antilles and 15 mya for Lesser Antilles (MacPhee & Iturralde-Vinent, 1994; Iturralde-Vinent 2006), it is still possible that other existing island chains facilitated dispersal (Iturralde-Vinent & MacPhee 1999). For instance, as it moved eastward, the Caribbean plate’s leading edge might have provided an island corridor — the proto-Greater Antilles — which allowed for dispersal (albeit probably limited) between Central America and South America during the Middle Eocene, approximately since 49-45 mya (Iturralde-Vinent and MacPhee 1999; Ali, 2012; Roncal, Nieto-Blázquez, Cardona & Bacon, 2020). Additionally, other proposed paleogeographical scenarios such as ‘GrANoLA’ — a Greater Antilles-Northern Lesser Antilles intra-oceanic subaerial connection (Philippon *et al*., 2020) — might also have played a role in the dispersal of dipsadid snakes from Central to South America, via continental islands (Cornee *et al*., 2021). Despite these ephemeral landmasses not being present in our analyses due to their disappearance (Iturralde-Vinent & MacPhee, 1999) and consequent lack of dipsadid records, jump dispersal likely played a role in the biogeographical history of this group, as supported by the +j (founder event) parameter in the best models.

Most lineages from the subfamily Xenodontinae diversified outside Central America and in the last million years in cis-Andean South America. One example is the tribe Alsophiini (Xenodontinae) which dispersed to and subsequently diversified in the West Indies during the Eocene-Oligocene transition (ca. 34 mya), which confirms that most of this insular extant fauna has a South American origin (Agnolin, Chimento & Lucero, 2019; Crews & Esposito, 2020), as previously suggested for Alsophiini (Hedges *et al*., 2009). This pattern and time frame are perfectly congruent with the GAARlandia scenario (Iturralde-Vinent & MacPhee, 1999). While the existence of GAARlandia has been increasingly questioned due to conflicting geological and paleo-oceanographic data (Ali, 2012; Ali & Hedges, 2021), several taxa with different dispersal abilities have been shown to have dispersed to the West Indies during this period such as giant sloths (Delsuc *et al*., 2019), arthropods (Crews & Esposito, 2020), freshwater fishes (Říčan, Piálek, Zardoya, Doadrio & Zrzavý, 2013), and amphisbaenids (Graboski *et al*., 2022). However, despite the congruent temporal window, it is still possible that West Indian xenodontines were the result of successive dispersal across the non-contiguous Aves Ridge, as suggested by the jump dispersal model and other taxa with similar patterns (Crews & Esposito, 2020, but see Ali & Hedges, 2021). Over-water dispersal seems to also be the process responsible for the more recent (∼ 10 mya) dispersal of *Erythrolamprus juliae* and *E. cursor* into the Lesser Antilles, since these islands are younger than 15 mya (Iturralde-Vinent, 2006), and thus long after GAARlandia had emerged and disappeared, as also shown for *Corallus* boids (Henderson & Hedges, 1995).

The timing of a contiguous land bridge, the Panama isthmus, between Central America and South America has been a hot debate topic among geologists, ecologists and biogeographers, with recent studies providing evidence that it likely occurred before the Late Miocene (∼ 10 mya) — much earlier than previously thought (∼ 3.5 mya; see Bacon *et al*., 2015; Buchs *et al*., 2019). While dipsadids entered South America before the earliest estimates of the formation of the Panama isthmus, there is evidence of recent expansion to and from Central America, coincident with other two significant increases in migration rate (Bacon *et al*., 2015). This expansion occurred mainly for dipsadines between 12 and 9 mya, with several genera (e.g., *Sibon* and *Imantodes*) reaching trans-Andean South America. This is also true for some species of Xenodontinae (e.g., *Oxyrhopus*, *Pseudoboa* and *Erythrolamprus* genera) which underwent the inverse path more recently — expanding from the cis-Andean region to the trans-Andean region and Central America. However, all these xenodontines are also broadly distributed in South America, and their presence is Central America represents the extreme north of their distributions. Besides Conophiini there is no species of Xenodontinae that is exclusive to Central America. Further studies might focus on the processes behind this pattern, especially if differences in phylogenetic niche conservatism for habitat or other ecological aspects might have played a role in this extension, as some species have marked habitat-associated distributions (Serrano, Vieira-Alencar, Díaz-Ricaurte & Nogueira, 2020; Serrano *et al*., 2023).

Regarding the Early Miocene of dipsadines from Central America into South America, different processes may be involved, as show for two closely related clades in close temporal proximity: representants of the tribe Dipsadini between 20-25, and the speciose genera *Geophis* and *Atractus* at around 22 mya. The ancestor of both theses clades was Central American, but our results suggest that the ancestor of Dipsadini first extended its distribution to South America and later underwent vicariance, while *Atractus* most likely jump dispersed. Even though there was no contiguous landmass connecting the two continents at that time, other proposed hypotheses might explain how these two clades entered present-day trans-Andean South America: stepping-stone dispersal by volcanic island chains and/or over-water dispersal, both facilitated by the collision of the Choco block with the South American continent (North Andean block; Bacon *et al*., 2015; Buchs *et al*., 2019) in the Early Miocene, at around 25-23 mya, corroborated by thermochronology and changes in geochemical profiles (Farris *et al*., 2011). Furthermore, this aligns with another significant increase in migration rates between the two continents (Bacon *et al*., 2015). While the exact timing for a contiguous terrestrial connection between Central America and South America is disputed (O’Dea *et al*., 2016, but see Jaramillo *et al*., 2017; Molnar, 2017), the formation of a land bridge is a complex and gradual process which might have allowed for over-water or stepping-stone dispersal into present-day trans-Andean South America over time, as suggested for other taxa (O’Dea *et al*., 2016), including dipsadid snakes of the genera *Imantodes* and *Leptodeira* (Daza *et al*., 2009; Costa *et al*., 2022).

The collision of the Choco and North Andean blocks in Early Miocene allowed for biotic dispersal between the two continental masses, and also triggered important geological changes in South America: increased Andean orogenesis and propagation of the Llanos basin (Farris *et al*., 2011; Mora *et al*., 2020). While exhumation of the Andean metamorphic rocks had likely began in the Late Cretaceous (before the Andean orogenesis, around ∼ 100 mya; Avellaneda-Jiménez *et al*., 2020), the uplift of its northernmost portions (e.g., the Central and Western Cordilleras) significantly accelerated in the Miocene, around 23 mya (Hoorn *et al*. 2010; Chen, Wu & Suppe, 2019). As a consequence, diversification increased for several plant and animal taxa and the Dipsadinae were no exception. Our results show that the early diversifications of the tribe Dipsadini and the genus *Atractus* in South America are congruent with peak uplifts in early Miocene (∼23 mya), similarly to Aromabatidae frogs (Boschman & Condamine, 2021) and clearwing butterflies (Elias *et al*., 2009), even though a large portion of the Andes was at half its present elevation (Gregory-Wodzicki, 2000). An increasing geographical and genetic isolation likely occurred for species with cross-Andean distributions imposed by Andean uplift that subsequently led to a pattern of coeval cis-Andean/trans-Andean vicariant events in Dipsadidae – within the *Atractus* genus at 11 mya, as previously suggested (Passos, Lynch & Fernandes, 2008) – and in Xenondontinae, in the *Siphlophis* genus (∼ 8 mya), as well as for Neotropical pitvipers (Pontes-Nogueira, Martins, Alencar & Sawaya, 2021). The Andean uplift may have indirectly contributed to speciation by altering climate and environment in pan-Amazonia (Hoorn *et al*., 2010), as such events have been shown to be strong drivers of diversification in the region (Pinto-Ledezma, Simon, Diniz-Filho & Villalobos, 2017; Rangel *et al*. 2018; Vasconcelos *et al*., 2020), especially for ectotherms (Santos, Coloma & Summers, 2009; Esquerré, Brennan, Catullo, Torres-Pérez & Keogh, 2019; Meseguer *et al*., 2021). Further intense pulses of Andean Mountain building in middle Miocene (∼12 mya) and early Pliocene (∼4.5 Ma) coincide with potential cis-Andean/trans-Andean dispersal in xenodontine clades (the tribes Pseudoboini and Tachymenini, and in the genus *Erythrolamprus*) as well as increased speciation in *Atractus*. These direct and indirect effects of mountain uplift corroborate the role of the Andes as a “species pump”, increasing species diversification into surrounding environments such as the Amazon and the Choco (Rangel *et al*., 2018, Rahbek *et al*., 2019).

However, some trans/cis Andean biogeographical patterns within Xenodontinae seem to have been only indirectly impacted by the uplift of the Andes. The ancestral of the genus *Pseudalsophis* jump-dispersed to trans-Andean region—and after to the Galapagos Archipelago (Zaher *et al*., 2018)—around 34 mya, long before the first stages of the Andean uplift. Similarly, *Saphenophis* and the recently described trans-Andean genus *Incaspis* (not sampled in our analysis) have been recovered as independent and old lineages (tribes Saphenophiini and Incaspidini). The latter is strongly recovered as sister to the cis-Andean tribes Tropidodryadini and Philodryadini (Arredondo *et al*., 2020), likely suggesting other dispersal events between both regions in the early Oligocene, before the mountain uplift.

Our results show that current biogeographical patterns of the family Dipsadidae, the most species rich snake clade in the Neotropical region, have been shaped by complex evolutionary and geological processes. Our reconstructed model recovered an Asian origin for the Dipsadidae family and potential significant paleogeographical events such as Eocene land bridges, Andean uplift and the formation of the Panama isthmus. While both dipsadines and xenodontines originated in Central America, they showed different evolutionary and biogeographical trajectories since they have dispersed into South America at different time periods and in two different regions: trans-Andean and cis-Andean South America. This is likely responsible for not only their present distribution, co-occurrence and regionalization patterns but also for relevant differences in their ecology and richness, which may help to explain why both these two Neotropical subfamilies are much richer than their Asian and North American counterparts (Cadle & Greene, 1983; Serrano *et al*., in prep). Additionally, our results provide a refinement on the understanding of the historical biogeography of the Neotropical region and how important events have shaped its biota.

## Supporting information

Supp. Mat. 1

Supp. Mat. 2

## ACKNOWLEDGMENTS

We thank Richard Grenyer, Marcio Martins, Fabricio Villalobos, Paulo Inácio Prado and Fernanda Werneck for their valuable comments and suggestions. FS and MPN was financed by the Coordenação de Aperfeiçoamento de Pessoal de Nível Superior (CAPES, Finance Code 001). RJS CN received funds from Fundação de Amparo à Pesquisa do Estado de São Paulo (Grant/Award Number: 2015-20215-7). RJS received funds from Fundação de Amparo à Pesquisa do Estado de São Paulo, Grant/Award Number: 2020/12658-4).

## DATA AVAILABILITY

Data are available from Figshare: to be added

## BIOSKETCH

Filipe C. Serrano is an ecologist interested in the spatial distribution of evolutionary processes, with a focus on herpetofauna. His current research topics include investigating how distribution at different scales may be used to infer ecological relationships and conservation status. This article represents the first chapter of his doctoral project “Phylogenetic diversity, endemism and conservation of cis-andean Dipsadid snakes” at the University of São Paulo (Brazil).

## AUTHOR CONTRIBUTIONS

FCS, MPN and FG conceived and designed the study. FCS, MPN and FG analysed the data. FCS wrote the paper. MPN, FG, CN, RJS and LRVA contributed critically to the drafts and all authors gave final approval for publication of the paper. This research has not previously been presented elsewhere.

